# Anchored Brownian motion and Bayesian methods for the analysis of single particle tracking data

**DOI:** 10.64898/2026.04.20.719631

**Authors:** Alan Boles, Robert J. Pyron, Sarah Shelby, Ioannis Sgouralis

## Abstract

We present a novel statistical method and a prototype computational implementation for estimating the diffusion coefficient from single particle tracking (SPT) data. Our method is based on anchored Brownian motion which is a novel representation that relaxes the restrictions of conventional Brownian motion. Our method is fully developed in Bayesian terms and allows for robust estimation of diffusion coefficient and quantification of the uncertainly propagated from limited data quantity and quality as appropriate for the analysis of live-cell SPT data. We compare our methods with conventional Brownian motion and demonstrate superior performance in estimating the correct value of the diffusion coefficient. Finally, we benchmark our methods with SPT data from in cellulo and in silico experiments.

## 1 Introduction

Ubiquitous in the natural and life sciences, diffusion governs the random movement of microscopic particles through a physical medium, such as a volume or a surface. The diffusion coefficient, characteristic of diffusive motion, controls the rate at which a molecule or particle moves through that medium and the overall time it takes to explore the region accessible to it [45, 55, 83]. The diffusion coefficient is an important feature and frequently a determining factor in the feasibility of empirical investigations in physics, chemistry, and biomedicine [2, 86]. Due to its importance, several families of analysis methods for the characterization of the diffusion coefficient from *raw* observation data are prevalent in the literature [20, 48, 84, 75, 5, 77, 61] and are actively investigated within physics, mathematics, and, more recently, machine learning [16, 70, 56, 11, 47, 80].

The prominent members of this family include bulk correlative methods such as fluorescence correlation spectroscopy (FCS) [17, 39] and fluorescence recovery after photobleaching (FRAP) [13, 68, 22]. These techniques [88, 32], and their variants [26, 23, 87, 39], probe particle dynamics indirectly through the variation of the fluorescence intensity emitted by labeled molecules that move randomly in bulk past a narrowly defined physical spot.

Another widely used type of analysis methods involves single-particle tracking (SPT) [33, 71, 74]. Most methods for estimating diffusion coefficients that rely on SPT use wide-field microscopes to collect image data that directly capture the random motion of individual particles in successive time instances [49]. Provided localizations obtained after processing raw experimentally acquired images, SPT-based methods proceed with data analysis through calculations on the extracted single-particle trajectories. In contrast to indirect data, SPT data record movement directly as paths of successive positions of individual particles [69, 85] and allows for a more detailed characterization of the diffusion coefficient than with bulk methods [75, 5, 77, 61].

The theoretical foundation of SPT data analysis is rooted in the mean squared displacement (MSD). For an ensemble of *M* particles moving in 1D, this is defined [42, 58, 46, 18] by the average

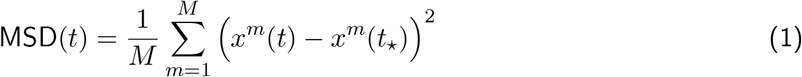

where superscripts *m* = 1, …, *M* label the particles in the ensemble and *x*^*m*^(*t*) denotes their position at time *t*. As can be seen, the MSD is a function of time *t* and this function is defined only up to an arbitrary but fixed reference time *t*_⋆_ that, in practice, is most often chosen to coincide with the time of the earliest available measurement or the onset of the experiment’s course. Given an MSD, a linear equation of the form

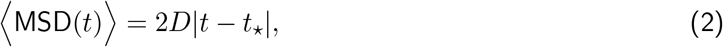

is invoked to relate the expectation of the MSD with the diffusion coefficient *D* under estimation [44, 42, 58]. In eq. (2), the expectation ⟨·⟩ is considered on the trajectories of the particles that make up the ensemble and serves to isolate random motion effects. Following SPT, eqs. (1) and (2) are combined to enable regression-type statistical methods for estimating *D*. For example, by plotting the empirical MSD at each time step of the available data and recovering *D* from the slope of the resulting graph using least squares optimization [42, 57, 4, 82, 8].

Despite its wide adoption, straightforward implementation, and low computational requirements, the results of MSD-mediated SPT-analyses are critically dependent on the quantity and quality of the data [78, 37]. Therefore, it is not surprising that they are susceptible to errors and biases [31, 41, 14, 43, 9]. Specifically, an empirical MSD relies on ensembles of a *finite number of particles* with each one of them probed for only a *finite number of steps* as only time-discrete versions of eqs. (1) and (2) can be formed and processed. In addition, an empirical MSD ignores observational noise and depends on the choice of reference time. Relying on finitely many discretized particle trajectories makes estimates of the diffusion coefficient susceptible to small-sample errors that, in the existing literature, are typically left unquantified [42, 84]. Ignoring the localization error without nontrivial changes in the definition of the MSD leads to biased estimates of the diffusion coefficient that are generally manifested as overestimation [42, 43]. Finally, choosing a reference time for the calculation of an empirical MSD via eq. (2) either reduces the available data by removing 1 localization per particle, since it reserves this localization for referencing, or artifactually forcing the positions of all trajectories in an ensemble to coincide at the chosen time even if this time is set to the distant past of all probed trajectories’ onset. Both are crucial limitations for SPT datasets where trajectories, even sufficiently many of them, are only probed for a few time instances such as 2 or 3 successive localizations, or are probed only for short periods of time such as typically found in *live-cell acquisition methods* employing super-resolution modalities [15, 33, 79]. In addition, an empirical MSD follows remarkably non-Gaussian statistics, indicating that least squares is an inappropriate training method for regression such as in eq. (2).

In order to develop an accurate and rigorous method for estimating the diffusion coefficient from SPT data that relaxes the aforementioned limitations, analysis methods are needed that: (i) are not limited by the size of the datasets available either in terms of trajectories or localizations within a trajectory; (ii) account for localization noise; (iii) analyze every localization along a given trajectory and across multiple trajectories of an ensemble symmetrically without requiring a choice of a fixed reference time; and (iv) incorporate the exact diffusion statistics in training without involving least squares or similar assumptions.

In this study, we present a comprehensive Bayesian formulation to estimate the diffusion coefficient from SPT data that meets these requirements. First, we develop and present a novel mathematical framework to represent diffusive motion, termed *anchored Brownian motion*, that relaxes the limitations of conventional Brownian motion necessitating the choice of reference time. Next, using our novel formulation as the basis, we develop a principled statistical model and an efficient computational implementation, which allow for robust estimation of the diffusion coefficient and quantification of the uncertainly propagated from limited data quantity and quality that include small dataset sizes and noise. Finally, we benchmark our methods with SPT data from in cellulo and in silico experiments.

## 2 Methods

In this section, we first present a new theoretical framework for modeling diffusive motion. Our framework extends existing approaches on Brownian motion and relaxes some of its critical assumptions for SPT. In subsequent sections, we show how our novel framework can be extended and used to estimate the diffusion coefficient from SPT data with realistic characteristics.

For clarity, initially, we consider only motion in 1D. Our notions are naturally extended to multiple Cartesian dimensions, and we do so subsequently.

### 2.1 Anchored Brownian motion

Anchored Brownian motion (ABM) is the mathematical foundation for our model. ABM with diffusion coefficient *D* anchored at time *T* to the point *X*, is denoted by

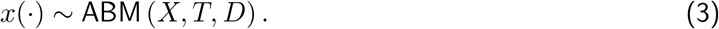

This is a random trajectory *x*(*t*), i.e. a stochastic process in the form of a random function [57, 51, 52], which is defined at all times *t*. By *anchor* we refer to time *T* and position *X*.

We define our ABM trajectory as a *Gaussian process* [60, 57] whose mean function depends on the position *X* and its covariance kernel depends on the time *T* and diffusion coefficient *D* according to

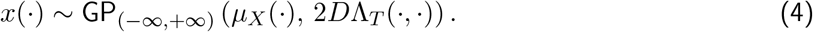

Specifically, in our definition of ABM the mean and covariance of the Gaussian process are determined by the functions

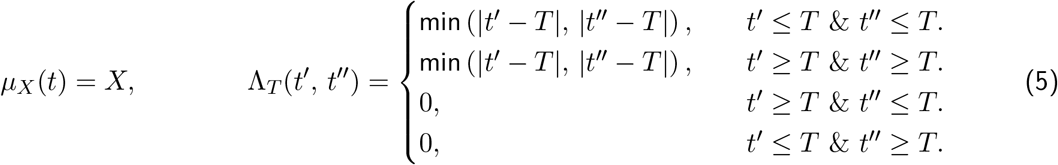

For comparison, we mention that the conventional Brownian motion (CBM) can also be defined as a Gaussian process [60, 52]. In this case,

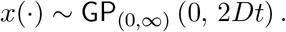

The critical differences [30, 62] between CBM and ABM include: (i) the time domain, with the former defined *only* for *t* ≥ 0; (ii) the placement of the reference position and time, with the former having it *fixed* at *X* = 0 and *T* = 0; and (iii) the trajectory’s position at *t* = 0, with the former *restricting* it to Figure S1 compares the two representations.

Extending the definition of CBM to ABM, we introduce an anchor with explicit position and time components into our framework for several reasons that are key to facilitating SPT data analysis. These include:

- Interpretation of SPT data and extraction of dynamical features require a motion model [33]. CBM assumes that the modeled trajectory passes through the spatiotemporal origin. Such an assumption makes interpretation dependent on the placement of the origin and this arbitrary choice imposes an asymmetry among the given localizations of a trajectory with those closer to the origin weighting more than those farther away. This choice biases *D*, as localizations closer to the origin have a stronger influence on the estimates than localizations farther away. With the inclusion of a general anchor, ABM makes this dependence explicit and parameterizes it in a physically meaningful manner. In this way, as we show later, we can incorporate uncertainty in the anchor, allowing it to float over both time and space, in a mathematically principled manner that maintains symmetry across space and time of all localizations in a trajectory.
- Additionally, our separate anchor allows all localizations along a given trajectory to be considered and analyzed similarly to the other ones, as none of them need to coincide with the anchor.
- Having a separate anchor that need not coincide with any of the localizations in a given trajectory, we are not only able to analyze all localizations in a dynamically equal setting, but also incorporate localization error with similar statistics across them.

Despite its more general formulation, ABM presents a valid model of Brownian motion dynamics appropriate for physical applications, since it still satisfies eq. (2). Specifically, considering an ensemble of anchored Brownian trajectories all of which share the same diffusion coefficient but, generally, different anchors, in the Supporting Information we show in detail that eq. (2) still holds. In addition, although not used directly in this study, we also derive the MSD’s exact statistics.

### 2.2 Data analysis with anchored Brownian motion

SPT data enables the characterization of the diffusion coefficient through successive position measurements along a particle’s trajectory. However, because the SPT data do not probe *D* directly, the values of *D* must be inferred by parameter estimation techniques. With the goal of inferring *D* from ABM position measurements, two mathematical problems are of practical interest: a direct and an inverse one [57, 76, 3].

Because SPT data are only available at discrete times [33, 15], here we focus on cases where an ABM trajectory *x*(·) is assessed only at given instances. For clarity, we consider *N* given time instances *t*_*n*_ that are indexed by subscripts *n* = 1, …, *N* and focus our presentation of the positions at these instances

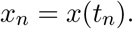

Without loss of generality, we consider only times that are arranged in increasing order *t*_*n*_ *< t*_*n*+1_.

In this context, our *direct problem* asks for a computational approach to the generation of ABM trajectories. Mathematically, this corresponds to random sampling from the distribution *p*(*x*_1:*N*_ |*X, T, D*). In contrast, the *inverse problem* asks to infer the ABM trajectory given noisy measurements [40]. That is, given noisy measurements, we seek to reconstruct ABM trajectories which may have given rise to these measurements. For clarity, we denote with *w*_*n*_ the position measurement made at time *t*_*n*_, and to proceed, we also adopt the noise representation given by

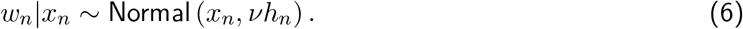

This generic model, which accounts for localization error, assumes additive Gaussian noise with variance *ν* that may differ between data points up to multiplicative factors *h*_*n*_. Mathematically, our inverse problem corresponds to a random sampling of the distribution *p*(*x*_1:*N*_ |*X, T, D, ν, w*_1:*N*_).

In the following, we present efficient algorithmic solutions to both problems. For simplicity, we assume that the parameters *X, T, D* and *ν* are fixed and attain known values. We relax this important assumption in subsequent sections, where we build upon the methods herein to develop a comprehensive Bayesian framework for SPT data analysis.

#### 2.2.1 Direct problem

Due to the definition of ABM as a Gaussian process [60, 57], sampling from *p*(*x*_1:*N*_ |*X, T, D*) consists of sampling a Gaussian vector, of size *N*, with mean and covariance function determined by eq. (5). The covariance matrix of this vector exhibits a block diagonal structure, with sizes of the blocks depending on the relative location of the times *t*_1:*N*_ to the ABM anchor *T*.

To be more precise, we denote with *P* and *Q* the last and first indexes *n* such that *t*_*n*_ *< T* and *t*_*n*_ *> T*, respectively. Under this convention, our Gaussian vector statistics take the form

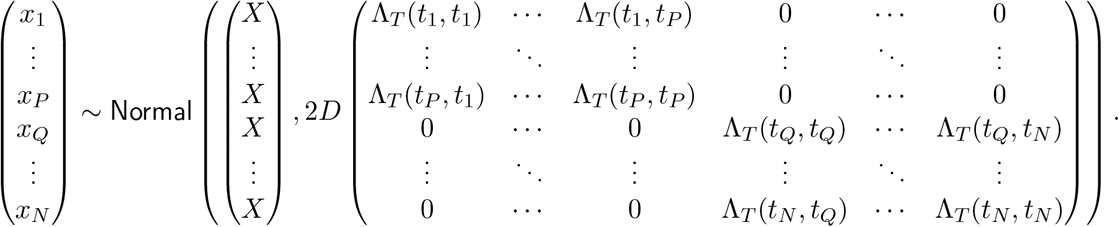

Because of the zero off-diagonal blocks in the covariance, this simplifies to two independent Gaussian vectors that encode the pre-anchor *x*_1:*P*_ and post-anchor *x*_*Q*:*N*_ parts of the trajectory separately. Therefore, for computational simplicity, sampling from *p*(*x*_1:*N*_ |*X, T, D*) proceeds by sampling separately a Gaussian vector with the pre-anchor and a Gaussian vector with the post-anchor positions. Taking advantage of the special form of the covariances within each diagonal block, both vectors can be sampled by generating univariate Gaussian random variables according to the following scheme

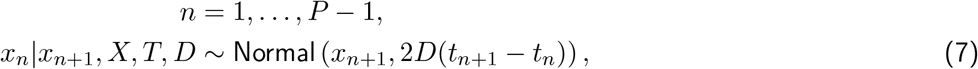

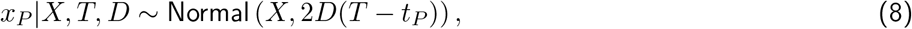

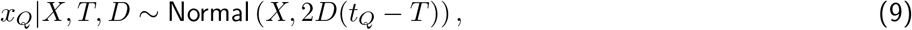

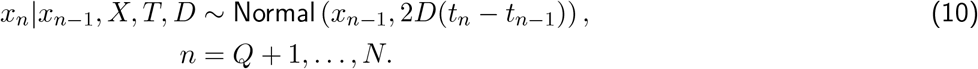

This scheme first samples the positions *x*_*P*_, *x*_*Q*_ closest to the anchor and proceeds *outward* through ancestral sampling [57, 34]. This univariate formulation makes sampling simpler and computationally efficient, as it avoids inversion of potentially large covariance matrices. Figure 1 (upper panel) demonstrates the direction of the computations.

**Figure 1:**
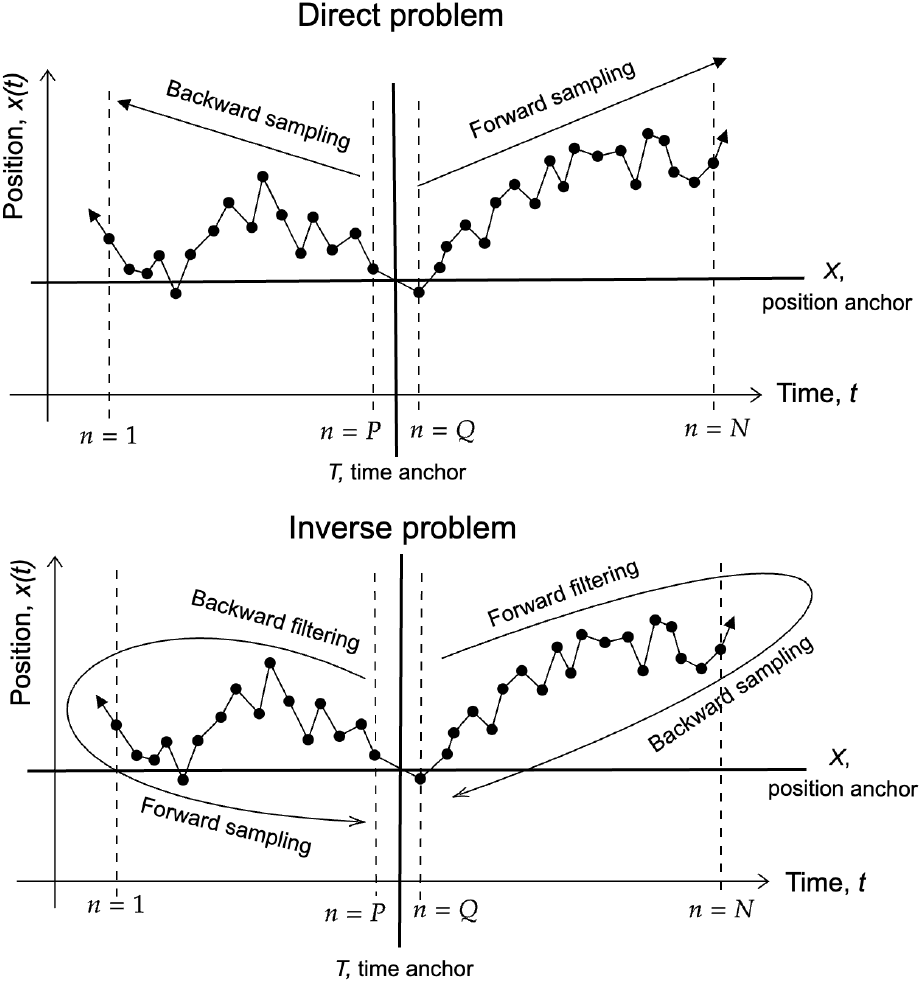
Sampling (top) and filtering (bottom) directions in the solution of the direct and inverse problem pertaining to anchored Brownian motion.

#### 2.2.2 Inverse problem

Our goal now is to sample from the probability distribution *p*(*x*_1:*N*_ |*X, T, D, ν, w*_1:*N*_) that, unlike before, conditions also on localization measurements *w*_1:*N*_. Due to the block-diagonal structure of the covariance and independence in the noise model of eq. (6), the sampling is based on the decomposition

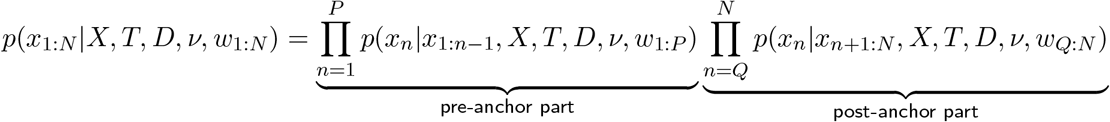

that keeps decoupled the pre- and post-anchor parts of the trajectory. Because every involved distribution in eqs. (6) to (10) is Gaussian and every position variable is linearly related to the others, each individual factor takes the form

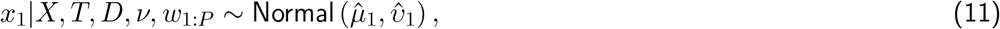

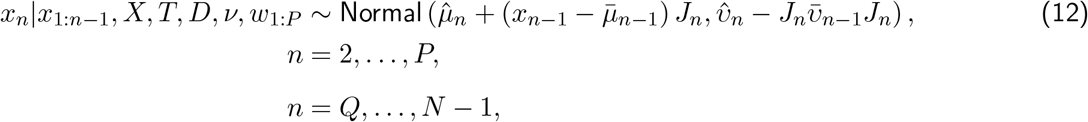

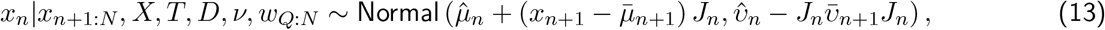

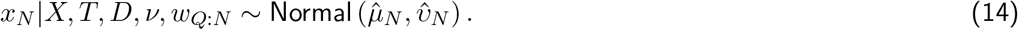

Consequently, starting from the outmost positions *x*_1_, *x*_*N*_ and moving *inward* toward the anchor, we perform ancestral sampling for the pre- and post-anchor parts separately. The statistics 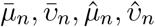 and *J*_*n*_, that must be computed in advance, are obtained using the formulas in the Supporting Information. These formulas proceed in the *opposite* direction, that is, start at the anchor and move outward. Figure 1 (lower panel) demonstrates the direction of the computations.

Essentially, our scheme is a modified version of Kalman filtering [28, 57, 29, 21] that accounts for the bidirectionality of the ABM motion as captured in eqs. (7) to (10). In addition to offering an elegant solution to our inverse problem, Kalman filtering is computationally efficient [25, 29, 21], and, also, with minor modifications, allows the estimation of the optimal trajectory by means of its smoothing counterparts [57, 21]. Since our main goal is to estimate *D*, we do not pursue this direction further in this study, but more details can be found in [57].

### 2.3 Bayesian single particle tracking data analysis

Having established our basic notions and algorithms, we now provide a general Bayesian formulation for the analysis of SPT data using ABM as our foundational representation of motion. To account for realistic SPT data, from now on we present our formulations in 2D. This includes 2D motion and 2D position data.

Our Bayesian model assimilates the information from successive noisy localizations of an ensemble of particle trajectories to arrive at an estimate for the ensemble’s diffusion coefficient. In our model, we consider trajectories of varying sizes and varying localization accuracy as obtained after processing raw experimentally acquired SPT images [15, 33].

#### 2.3.1 Bayesian model formulation

In the Bayesian context [57, 33, 81], our ABM model forms the prior on the trajectories, which, due to localization error, remain hidden. Because not only the diffusion coefficient but also the anchors are among the unknowns influencing our ABM trajectories, we assign hyperpriors to both its position and time. In this way, we allow our ABM to “float” across space and time, giving rise to a very flexible model to match a given trajectory’s data that, with the appropriate choices, maintain temporal symmetry among its localizations. Finally, because the localization accuracy is most commonly left uncharacterized by most SPT image processing methods [15, 33], we also model it as a random variable and assign an appropriate prior. This way, our model combines uncertainty on the actual particle positions, their spatiotemporal anchors, and their motion dynamics, and allows a self-consistent propagation of this uncertainty to the resulting estimates of the diffusion coefficient.

##### Model description

The following is a complete description of our Bayesian formulation, and fig. 2 shows its graphical representation.

**Figure 2:**
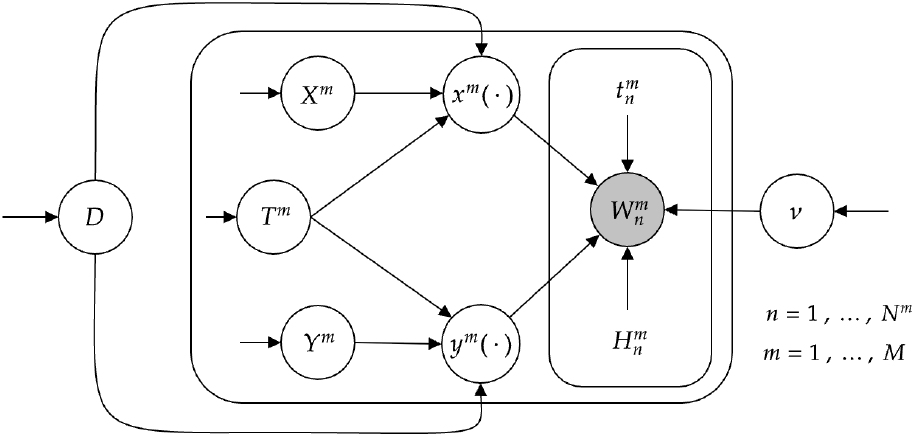
Graphical representation of our Bayesian model for SPT data analysis. Following the common convention [6, 57], random quantities are depicted within circles; while, fixed quantities are left free. Arrows indicate dependencies and plates repetition. Quantities with unknown values are left open and quantities with known values are shaded.

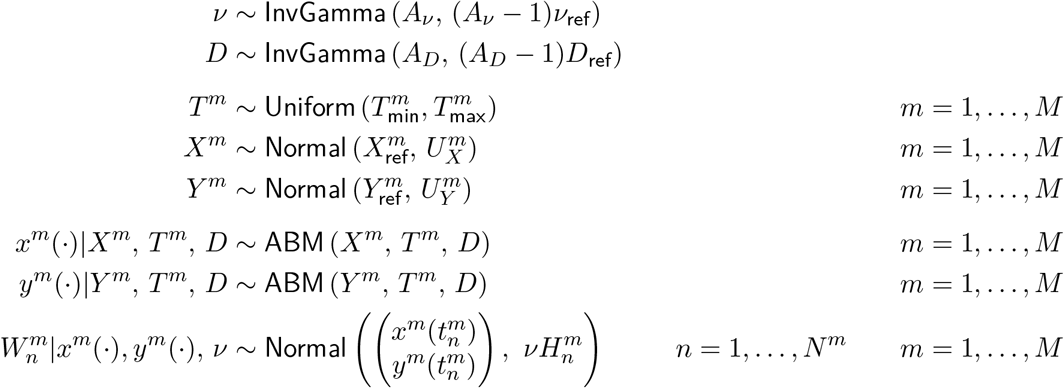

With this model, we consider an ensemble of *M* particles that diffuse in 2D that are denoted with superscripts *m* = 1, …, *M*. Each trajectory consists of its own successive localizations that we label with subscripts *n* = 1, …, *N* ^*m*^.

##### SPT data

In our model, the SPT data for each particle consist of 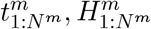 and 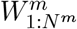. These model noisy assessments of successive positions of individual particles in our ensemble. In detail, 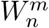 is the measurement of the *m*^th^ particle’s position at its *n*^th^ assessment and 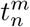 is its assessment time. To account for localizations in 2D, we employ the following notation

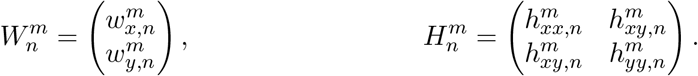

Here, 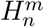 is a unitless square, symmetric, positive definite matrix that quantifies the localization accuracy in the two Cartesian coordinates and its correlation. In cases where image processing leading to SPT data yields no accuracy estimates, our 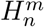 are set equal to the identity matrix.

##### Model variables

In our model, each particle is associated with two Cartesian trajectories *x*^*m*^(·), *y*^*m*^(·) and has an anchor determined by a time *T* ^*m*^ and a 2D position (*X*^*m*^, *Y* ^*m*^). Finally, *ν* and *D* are parameters common to all localizations and particles.

To complete our model, we choose the inverse Gamma prior distributions on *D* and *ν* to exploit conditional conjugacy [59]. Similarly, we choose Gaussian prior distributions on *X*^*m*^ and *Y* ^*m*^, again, to exploit conditional conjugacy. Regarding *T* ^*m*^, we choose a uniform distribution to induce a noninformative, but proper, prior and allow the time anchor to float over the data without a preference for particular time instances.

##### Posterior distribution

The joint posterior of our Bayesian model provides a probability distribution for each of the unknowns that conditions on the provided data. Discretizing each ABM trajectory *x*^*m*^(·), *y*^*m*^(·) over their respective times 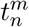 as in section 2.2, our joint posterior attains a probability density of the form

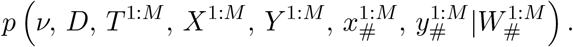

In this density, 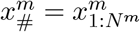 and 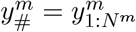 gather the discretized positions and 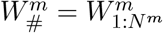 gather the observations of the *m*^th^ particle.

The form of our model’s posterior does not allow for closed formulas or direct sampling. For this reason, next, we develop a specialized Markov chain Monte Carlo sampler that can be used to provide estimates of the desired unknowns.

#### 2.3.2 Markov chain Monte Carlo

Markov chain Monte Carlo (MCMC) allows sampling from non-standard high-dimensional probability distributions [12, 34] such as our posterior. In particular, Gibbs sampling [19] is a prominent MCMC technique used to sample multivariate probability distributions in which random variables attain a natural grouping with simple conditional distributions that exploit conjugacy and graphical representations [59, 53, 57, 6, 63]. Because of these features, we develop a Gibbs sampling scheme that proceeds as follows:

- Initialize
  - generate *ν* by sampling from its prior
  - generate *D* by sampling from its prior
  - generate *T* ^1:*M*^, *X*^1:*M*^, *Y* ^1:*M*^ for each particle separately by sampling from their priors
  - generate 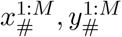 for each particle separately using the full conditionals via a 2D extension of the algorithm in section 2.2.2
- Iterate, in random order, the following stages until the requested number of samples is reached
  - update *ν* by sampling from its full conditional
  - update *D* by sampling from its full conditional
  - update *T* ^1:*M*^, *X*^1:*M*^, *Y* ^1:*M*^ by sampling from its full conditional for each particle separately
  - generate 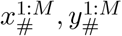 for each particle separately using the full conditionals via a 2D extension of the algorithm in section 2.2.2

Our Gibbs sampler and its implementation are described in detail in the Supporting Information.

To facilitate the adoption of our methods by practitioners, we also provide a fully working prototype implementation in our GitHub repository [7].

### 2.4 Data acquisition

#### 2.4.1 In cellulo data

##### Experimental description

For live-cell SPT experiments, the cell membrane proteins tracked were fusion constructs containing the photoconvertible fluorescent protein mEos3.2 fused to a minimal membrane insertion motif. Two constructs were used: one, termed “Src15-mEos3.2”, encodes the first 15 residues of the N-terminal myristoyl tail from the protein Src followed by a C-terminal mEos3.2. The second construct, termed “mEos3.2-trLAT”, encodes an extracellular N-terminal mEos3.2 linked to the single-pass transmembrane domain of the LAT protein, with the intracellular domain truncated after 10 residues. These constructs, including sequence and source information, are reported in [73].

Jurkat T cells were acquired from the American Type Culture Collection (ATCC) and maintained in culture at 37^◦^C with 5% CO_2_, in standard culture medium containing RPMI, 10% fetal bovine serum and 1% penicillin-streptomycin. For the expression of the mEos3.2 fusion constructs, cells were transiently transfected with plasmid DNA encoding mEos3.2-tagged membrane anchor constructs using a Neon XT electroporation apparatus (Thermofisher). 400-500×10^5^ cells were transfected with 1 *µ*g of DNA, then transferred to a cell culture flask containing equilibrated growth media and allowed to recover for 18-24 hours. MatTek culture dishes with glass coverslip bottoms were coated with poly-l-lysine (PLL) to adhere cells to the glass surface. For imaging, cells were transferred to PLL-coated dishes and allowed to adhere for 5 min at 37^◦^C. Before imaging, samples were rinsed with live-cell imaging buffer consisting of HEPES-buffered salt solution with BSA (HBSS-BSA: 30 mM HEPES, 5.6 mM glucose, 100 mM NaCl, 5 mM KCl, 1 mM KCl, 1 mM MgCl_2_, 1.8 mM CaCl_2_, 0.1% w/v BSA, pH 7.4).

Live-cell imaging was performed at room temperature on a Mad City Labs RM21 Advanced fluorescence microscope outfitted with a Total Internal Reflection Fluorescence (TIRF) illumination module, a TIRFLock active focal drift correction module, and an Olympus UPLAN APO 60x TIRF objective (NA = 1.50). The photoconvertible fluorescent protein mEos3.2 converts from green to red emission upon irradiation with 405nm light. To image single molecule trajectories of mEos3.2-tagged proteins, photoconversion of a sparse density of fluorophores was accomplished using low power illuminiation with a 405nm solid state laser (Coherent OBIS LX 405nm 50mW) and simultaneous excitation of the red-converted form with a 561 nm solid state laser (Coherent OBIS LS 561nm 150mW). Laser intensities were adjusted to maintain a low mEos3.2 photoconversion rate such that single mEos3.2 fluorophores were well separated from each other and image segmentation was unambiguous. Laser powers were generally between 5-10 kW/cm^2^ for the 561 nm laser, and 100-200 W/cm^2^ for the 405 nm laser. Simultaneous illumination with 405 and 561 nm lasers was accomplished using a quad-band dichroic mirror designed to reflect 405, 488, 561, and 647 nm lasers (Chroma ZT405/488/561/647rpc). Fluorescence emission was split using a Mad City Labs MadView multi-channel imaging system, with mEos3.2 red emission filtered through Chroma dichroic (Chroma T565lpxr) and emission (Chroma ET605/52m) filters. Images were captured using Micro-manager camera control software on a Hamamatsu ORCA-Quest qCMOS camera with CoaXpress fast image transfer. Integration times were maintained at 20 ms. Typically, ≈ 5000 total frames were acquired in 10 sec sequences of 500 frames that were saved as multi-page tiff files for further processing. 3-4 fields of view were collected of cells expressing both Src15-mEos3.2 and mEos3.2-trLAT.

##### Calibration measurements

A series of flat images were also acquired to calibrate the gain and variance parameters of individual pixels from the qCMOS camera, as previously described [71, 24]. Briefly, pixel offset and variance are calculated from a set of “dark” images acquired in the absence of signal, and gain is calculated from sets of “bright” images acquired at a series of increasing illumination intensity levels from a diffuse, uniform light source. 60,000 dark images were acquired with the camera lens cap in place, and 2730 bright images were then acquired at 10 set intensity levels. For this, a white LED source was placed ≈ 50 cm from the camera chip. A lens tube containing a ground glass diffuser disk was attached to the camera c-mount, and for each light intensity level, additional ND filters were inserted into the lens tube. For calibration images, acquisition settings including pixel binning and exposure times were as in the live-cell experiments.

##### Image processing

Once the raw image data are acquired, the images need to be processed before the SPT data are properly formatted for the ABM model of section 2.3.1. the The Supporting Information provides a schematic outline of our processing pipeline starting with raw tiff images. Our image processing pipeline has three main stages: camera calibration, image segmentation, and particle localization. In camera calibration, we use the calibration data and derive pixel maps for camera read-out offset, variance, and gain as described in [71]. In image segmentation, tiff images are displayed, and diffraction-limited spots are selected in each image by visual inspection. The pixel information is stored temporarily, as well as the gains, offsets, and variances necessary for localization. This process is carried out for each of the images until the desired amount of data is acquired. The data is then processed using a Bayesian localization method described in [71]. This method provides estimated locations of the particles, as well as covariance matrices, as required by the ABM model. The data output from this localization method corresponds to 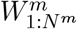 and 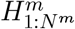. The times 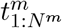 are calculated during the segmentation process by the start times of each frame’s integration.

Finally, the extracted SPT data are stored in formatted txt files which are, in turn, passed to the MCMC sampler of section 2.3.2, which outputs the estimated posterior distribution of the diffusion coefficient.

#### 2.4.2 In silico data

To generate synthetic SPT data, we used computer simulations. For these simulations, we prescribed values for *D, ν, M*, that mimic live-cell conditions. The sizes of the trajectories, *N* ^1:*M*^, are randomly selected using a geometric distribution with a prescribed mean, so we gain control of their average. We generate the corresponding SPT data by simulating the model of section 2.3.1 and generating the trajectories through a 2D extension of the algorithm in section 2.2.1. All simulations require common pseudorandom procedures, such as generating random variates, which we carried out with standard programming commands. For all simulations, the prescribed values were retained and used as “ground truth” values for comparisons with the results. The prescribed values were not used otherwise.

## 3 Results

We present a new method for analyzing SPT data and estimating diffusion coefficients. Our data consist of empirical assessments of successive positions along the trajectories of a particle ensemble. Such data may contain a varying number of localizations per particle or particles per ensemble. In addition, each localization can be contaminated with noise and may differ from the actual particle position, as is typical for SPT data obtained via image processing. Our method is fully developed in Bayesian terms and so provides not only estimates of the diffusion coefficient but also quantifies errors that arise from small datasets or propagated due to noise. Although our methods are compatible with MSD-based results, our analysis requires the construction of no MSD. Instead, it is based on a novel representation of Brownian motion that relaxes several of the key limitations of MSD.

In order to show how our method is applied and demonstrate its validity, we analyze experimental SPT data obtained using live-cell super-resolution fluorescence microscopy and qualitatively similar synthetic SPT data. With experimental data, we focus on two cases of characteristically different diffusion coefficients [73] and demonstrate that our method not only distinguishes between them, but also yields estimates in agreement with the existing literature. With the synthetic data, we compare the performance of our model directly against the ground truth and demonstrate that our method identifies the correct values in broad, experimentally relevant settings. We also compare our method to traditional methods that rely on CBM and demonstrate that our novel representation that relies on ABM outperforms them. Finally, we use our model to provide insight into the design and configuration of SPT experiments optimized for the estimation of diffusion coefficients.

### 3.1 In cellulo data

To test our method, we conducted separate laboratory experiments by imaging two different plasma membrane proteins in live cells. Both proteins are minimal membrane anchor motifs fused to the photoconvertible fluorescent protein mEos3.2. Our proteins have different structures and exhibit different diffusion behaviors [73]. Namely, Src15-mEos3.2 encodes the membrane anchor motif of the protein Src and consists of a post-translationally attached lipid anchor and a short sequence of positively charged amino acids, followed by mEos3.2; while, mEos3.2-trLAT is tagged at the N terminus with an extracellular mEos3.2, followed by the membrane anchorage motif consisting of the single transmembrane alpha helix from the protein Linker for Activation of T cells (LAT). Previous reports [73] have shown that mEos3.2-trLAT diffuses slower in cellular plasma membranes than Src15-mEos3.2.

Once we process the raw images acquired in the experiments and extract SPT data, we apply our ABM model to two datasets of ≈ 30 trajectories containing ≈ 100 displacements each. By maintaining the same number of trajectories and total displacements, *M* and ∑_*m*_ *N* ^*m*^ respectively, in both experiments, we ensure that our method is equipped with the same data volume and therefore any resulting differences are due to the dynamical characteristics of the two specimens rather than mismatched sample sizes.

Figures 3 and 4 summarize the data and present the estimated posterior distributions of the diffusion coefficients for the two experiments. With MAP estimates at 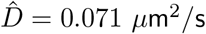 and 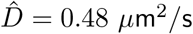, respectively, our analyses agree with previous reports in [73] on the diffusion rates of the two molecules. This shows that our ABM model is capable of distinguishing between different diffusion rates when applied to laboratory in cellulo data and also that the estimates it provides are compatible with existing studies.

**Figure 3:**
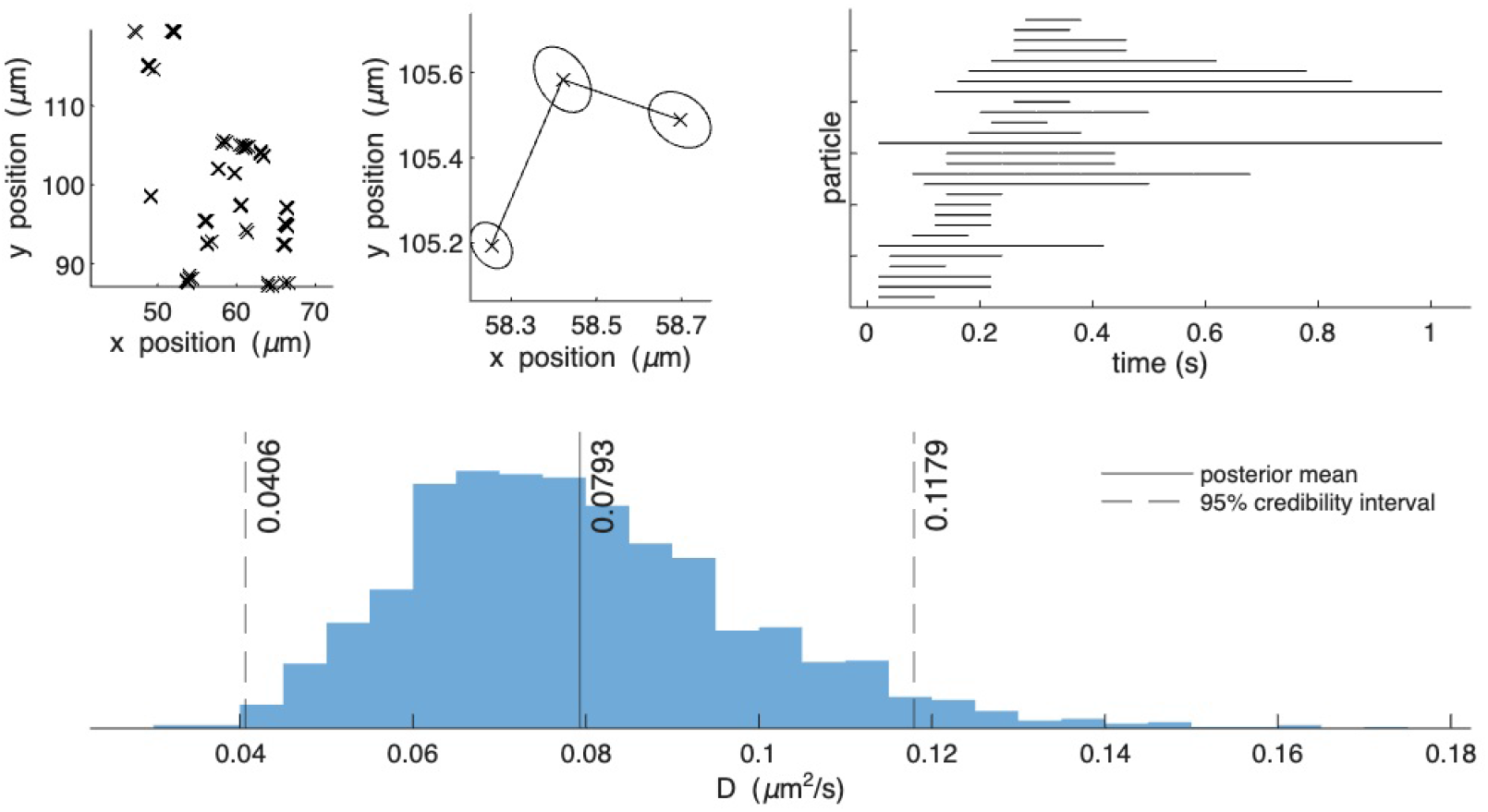
Summary of the mEos3.2-trLAT data set, along with the estimated posterior distribution for the diffusion coefficient. Top-left: spatial distribution of the localizations of all trajectories within the field of view. Top-middle: a representative path illustrating typical step sizes with ellipses indicating the localization errors. Top-right: temporal distribution of the localizations of all trajectories. Bottom: inferred posterior probability density for the diffusion coefficient D obtained from our ABM model; the posterior mean and the 95% credible interval are indicated with vertical lines.

**Figure 4:**
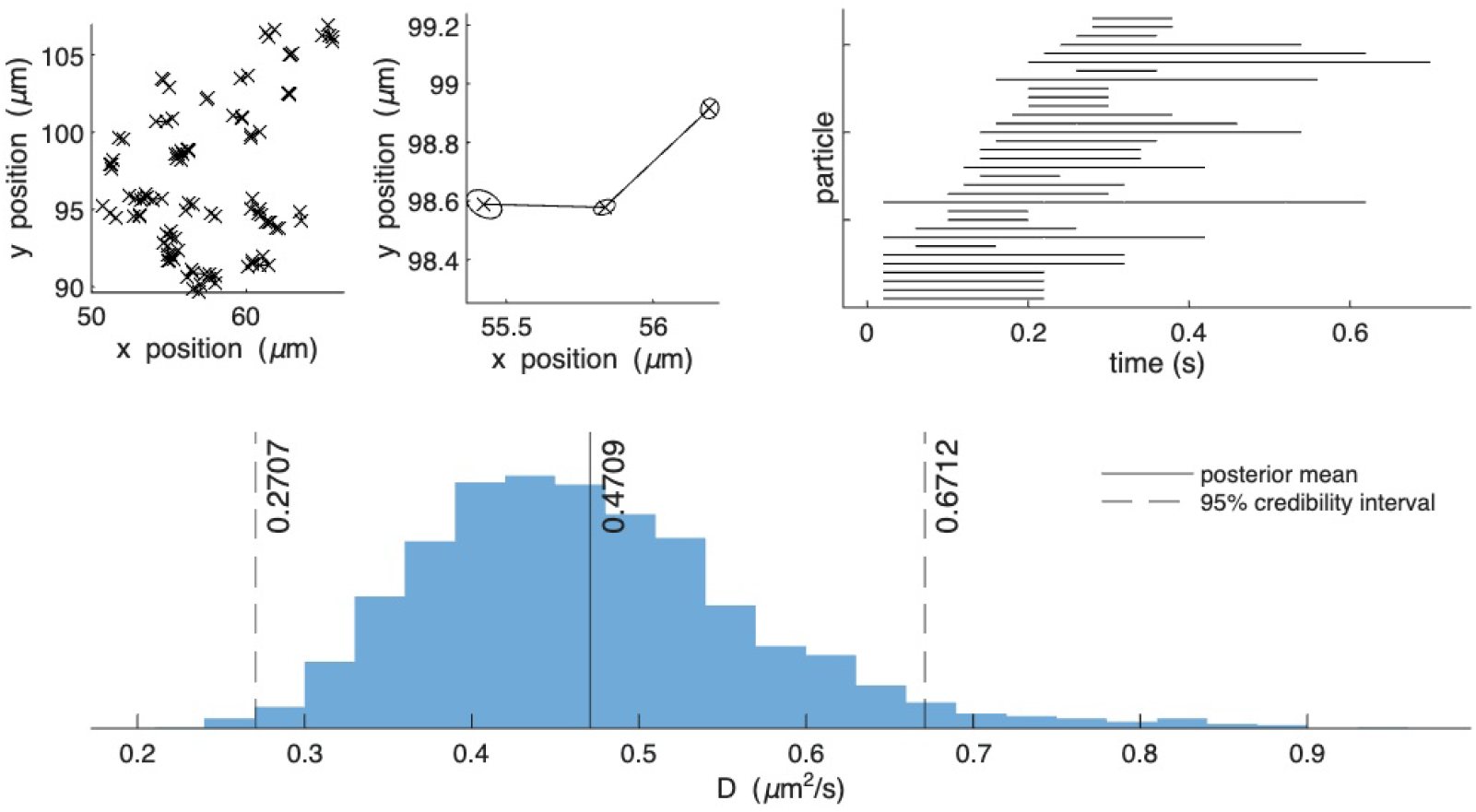
Summary of the Src15-mEos3.2 data set, along with the estimated posterior distribution for the diffusion coefficient. Top-left: spatial distribution of the localizations of all trajectories within the field of view. Top-middle: a representative path illustrating typical step sizes with ellipses indicating the localization errors. Top-right: temporal distribution of the localizations of all trajectories. Bottom: inferred posterior probability density for the diffusion coefficient D obtained from our ABM model; the posterior mean and the 95% credible interval are indicated with vertical lines.

### 3.2 In silico data

We now assess our method’s performance under a wider range of conditions that extend beyond our conducted experiments. For this task, we use synthetic SPT data, so we benchmark directly against the ground truth.

To verify that our estimates are indeed in agreement with the ground truth, we generate data with prescribed *D* = 0.10 *µ*m^2^/s and *ν* = 0.025 *µ*m^2^. As before, we generate 30 trajectories with ≈3 − 5 localizations each, for a total of ≈100 displacements. Our configuration mimics the livecell imaging data shown in fig. 3 obtained with mEos3.2-trLAT. Figure 5 summarizes our data and presents the estimated posterior distributions of the diffusion coefficient. With a MAP estimate at 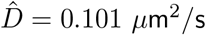, our analysis agrees remarkably well with the ground truth *D*.

**Figure 5:**
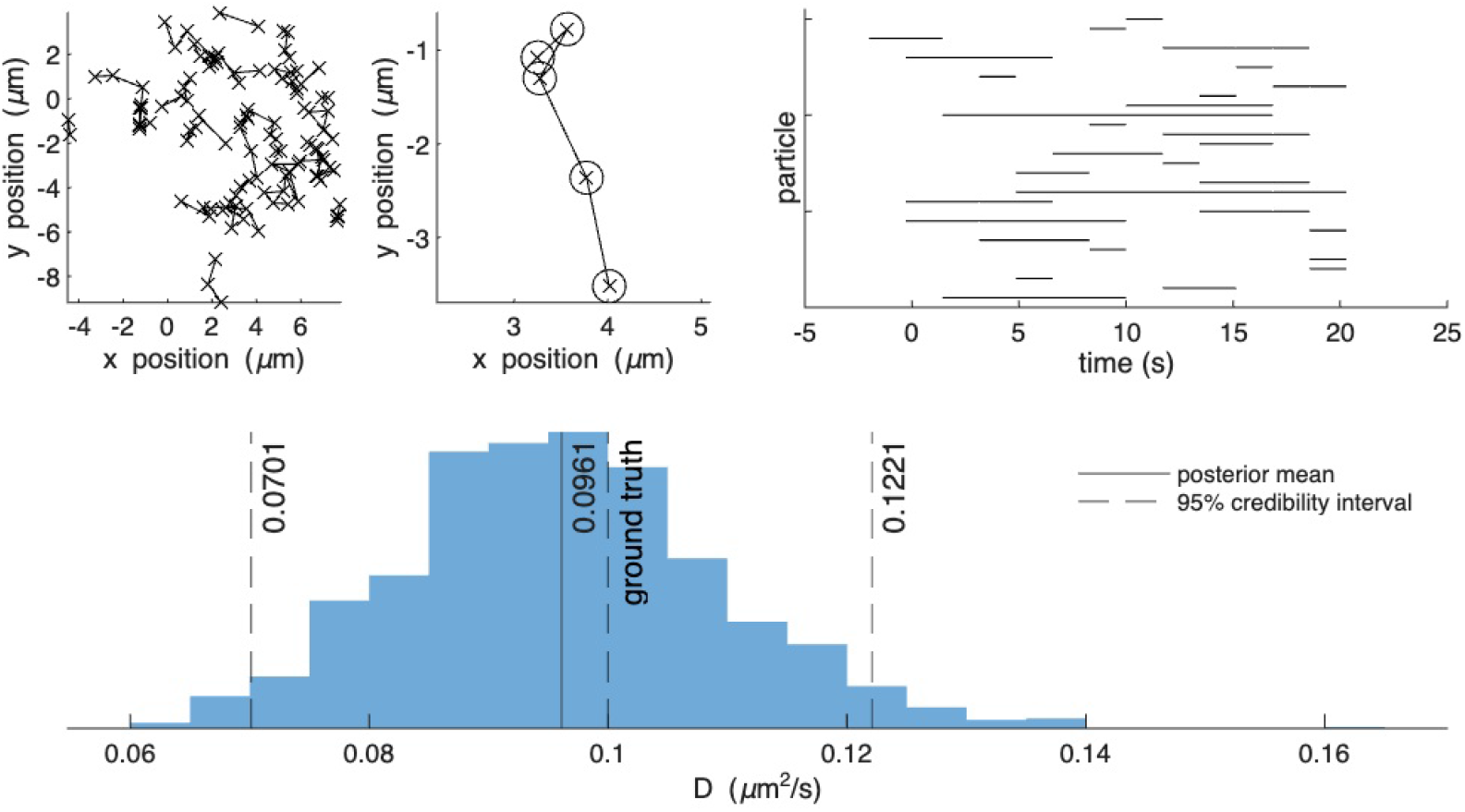
Summary of a synthetic data set mimicking the data of fig. 3, along with the estimated posterior distribution for the diffusion coefficient. Top-left: spatial distribution of the localizations of all trajectories within the field of view. Top-middle: a representative path illustrating typical step sizes with ellipses indicating the localization errors. Top-right: temporal distribution of the localizations of all trajectories. Bottom: inferred posterior probability density for the diffusion coefficient D obtained from our ABM model; the posterior mean and the 95% credible interval are indicated with vertical lines.

Using synthetic data, we also determine the effect of relying on ABM and its advantages over CBM for the estimation of *D*. Specifically, to test the effect of the anchor, we perform two different analyses on the same synthetic data. For both analyses, we use the same Gibbs sampler. However, in one case, we restrict the time of the anchor to coincide with the time of the first data point for each of the *M* trajectories. Similarly, we restrict the position of the anchor to coincide with the position of the first data point. Our modifications result in effectively a Bayesian method for estimating the diffusion coefficient under CBM. In the other case, our sampler is unmodified and estimates the diffusion coefficient with ABM.

Figure 6 summarizes the results of the two analyses. This figure compares the posterior estimates of the two cases for increasing data volumes as determined by the total number of displacements (ie. ∑_*m*_ *N* ^*m*^) in each ensemble. Our plots also highlight the means and 95% credibility intervals of the posterior estimates, along with the ground-truth *D*. As can be seen, for small datasets, ie. ≈ 200 displacements or less, both cases result in vague posteriors that peak away from the ground truth and spread over a considerably wide range of possible *D* values. In this regime, CBM estimates are associated with smaller credible intervals than ABM, indicating misleading confidence in the wrong values. Nevertheless, for larger datasets, ie. ≈ 1000 displacements or more, CBM consistently overestimates *D*. In contrast, ABM is able to converge to ground truth. This indicates that the inculcation of an anchor that is free to adapt to the data (as in the ABM model) leads to accurate and unbiased estimates; however, the inculcation of a fixed anchor (CBM model) leads to biased estimates with the bias persisting irrespective of data volume.

**Figure 6:**
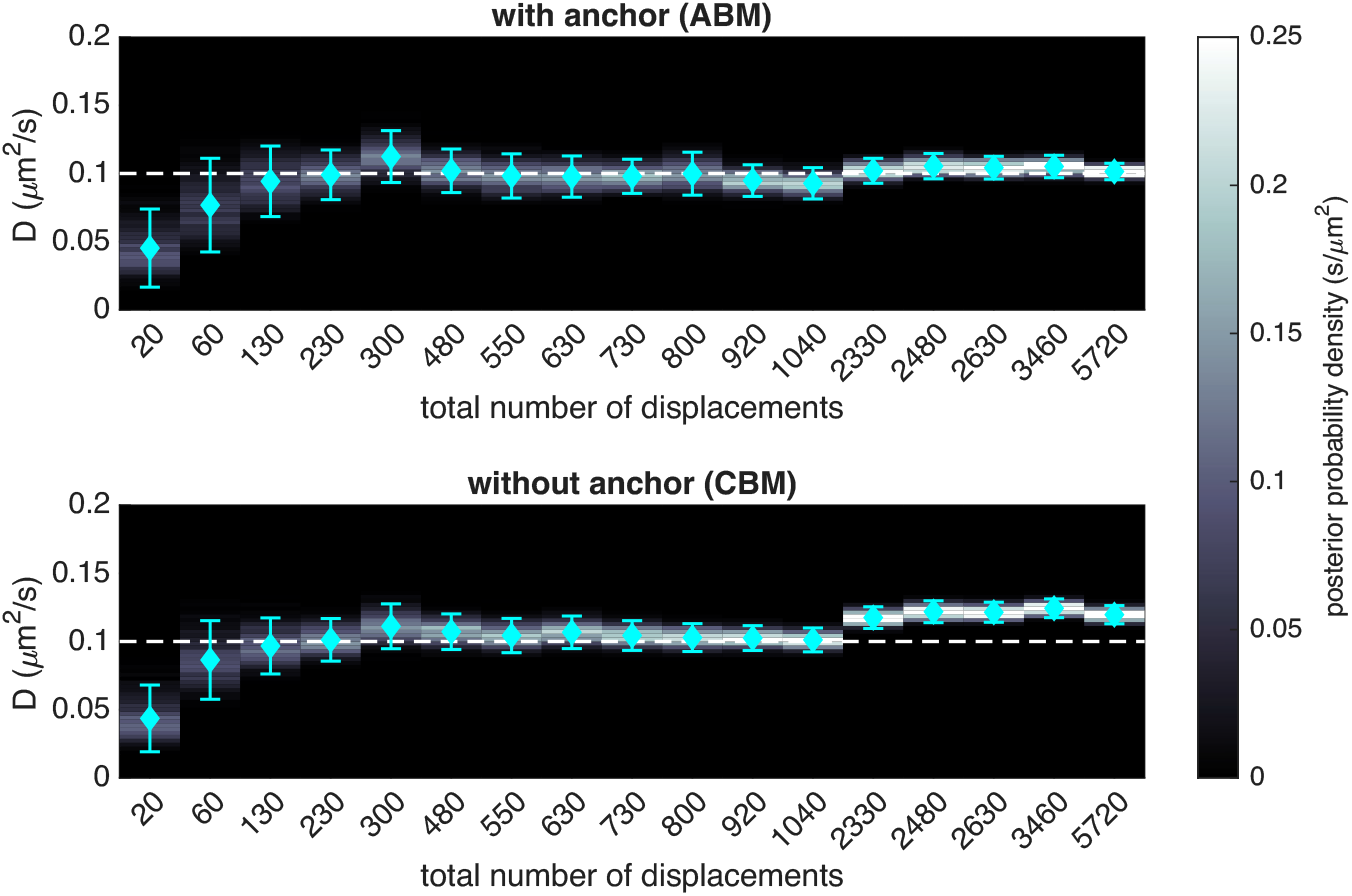
Comparison of ABM (top) and CBM (bottom) estimation performance across simulated conditions, highlighting accuracy and uncertainty quantification at increasing SPT data volume. For both panels the posterior distributions are color-coded and their mean and 95% credible intervals indicated with error-bars. The ground truth value is also shown with a horizontal line. At larger data volumes both ABM and CBM result in narrower credible intervals; however, only ABM converges to the ground truth, while CBM remains biased.

In addition to the estimation of diffusion coefficients, our novel methods, which allow both the generation and analysis of synthetic SPT data under prescribed conditions, can be used to guide the configuration of SPT experiments. This is particularly relevant for the configuration of super-resolution experiments, where, by adjusting particle density and illumination levels, practitioners need to balance imaging conditions leading to multiple short trajectories or fewer longer ones [36, 54].

To investigate which case might be preferable for optimizing the estimation of diffusion coefficients, we consider a simplified scenario. Specifically, we consider imaging conditions similar to section 2.4.2, and generate multiple SPT datasets of varying size. We constrain all datasets to 500 displacements each. However, we try cases where these displacements are distributed among a few excessively long trajectories or among multiple short ones. At one extreme end, we obtain a dataset containing 1 particle with 500 jumps. At the other extreme, we obtain a dataset containing 120 particles with 3-5 displacements each. We fill in the cases between these two extremes with more moderately evenly distributed SPT data, with the most average one containing 50 particles, with around 10 displacements per trajectory.

Figure 7 displays the mean and standard deviation of the estimated posterior distributions for the diffusion coefficient resulting from our ABM model for each of these cases. As can be seen, our results indicate that, although the posterior means are relatively stable across all cases with minor fluctuation around the ground truth, the standard deviation exhibits a decreasing trend at datasets of more trajectories despite the fact that these trajectories are shorter. Accordingly, our results support a preference for SPT datasets with more but short trajectories rather than a few longer ones, as the former are associated with narrower confidence intervals.

**Figure 7:**
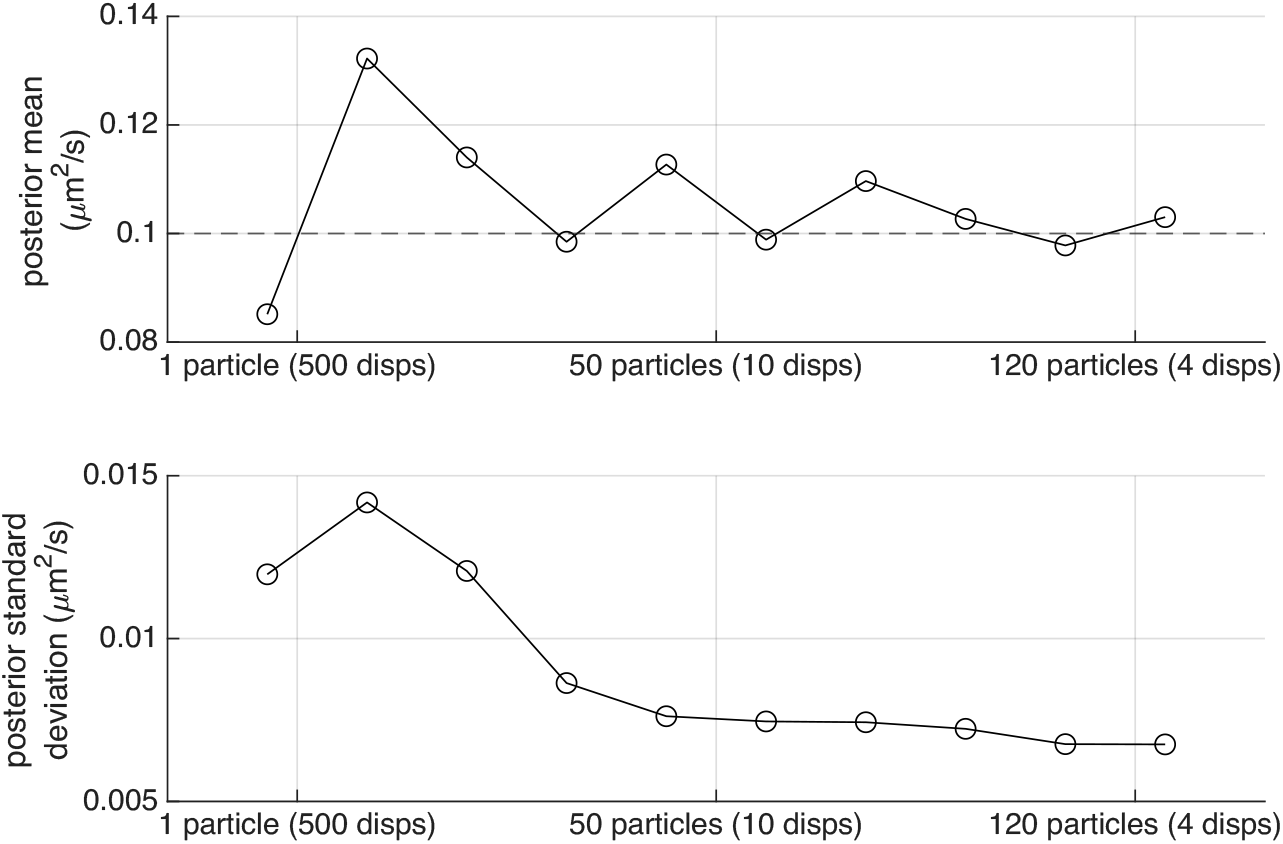
A comparison of posterior mean and standard deviation for estimates of the diffusion coefficient D across different amounts of localization data. From left to right: increasing number of particles and decreasing number of displacements per particle. For all posteriors shown here the total number of displacements over all particles combined is kept fixed at 500.

## 4 Discussion

The growing accessibility of single molecule techniques and the widespread adoption of super-resolution microscopy in biophysics and biochemistry research has lead to an abundance of information-rich data [10, 1, 35, 33, 15]. Leveraging such data in full requires new analysis methods that exceed in sophistication traditional approaches. With regard to diffusion models, especially for live-cell imaging applications, existing methods such as those mediated by empirical MSD curves rely on simplifying assumptions that are inappropriate or suboptimal for the analysis of experimentally acquired SPT data [33]. In particular, MSD-mediated analyses perform particularly poorly when SPT data stem from short trajectories or trajectories contaminated with localization error [43] which are common characteristics of live-cell SPT experiments.

In this study, we present a novel statistical methodology that addresses the main limitations in estimating diffusion coefficients from SPT data. In particular, we present a comprehensive Bayesian model which combines position data from multiple trajectories of arbitrary sizes. Our method explicitly accounts for localization errors provided by modern image processing workflows for super-resolution [71, 72, 50]. Our model relies on ABM which is a new approach to modeling Brownian motion that we develop herein. This representation of motion uses time reversibility to relax the fixed initial condition requirement of CBM. This allows a uniform prior distribution to be applied on the anchor of each trajectory, allowing objective data analysis that is not influenced by the choice of the anchor.

The validity of our approach is tested with synthetic SPT data as well as laboratory SPT data from live-cell single molecule tracking experiments. Characteristically, our methods lead to unbiased posterior estimates and allow for experiment design procedures that can help optimize imaging conditions.

However, despite its advantages for SPT data analysis, our approach also maintains drawbacks. For example, our approach relies on the ABM motion model, which is more complex than traditional approaches that rely on the simpler CBM motion model, leading to higher computational cost. For this reason, we developed specialized algorithms for its use. Additionally, the mathematical overhead for our approach is higher than that of MSD-mediated methods. These are limitations that pose barriers for practitioners. To facilitate a wider adoption, we provide a fully functional prototype implementation of our methods in [7]. Our implementation can be used as a stand-alone analysis application for SPT data and also as a component in larger workflows that incorporate photo-blinking [38, 64] and multi-color [65, 66, 67] or non-widefield [27, 39] imaging.

## Author contributions

A.B. contributed to the analysis methods, computational implementation, and software development. A.B. and I.S. prepared the manuscript. R.P. and S.S. provided the experimental data and commented on the manuscript. I.S. designed the research and supervised all aspects of the project.

---

**Figure S1:**
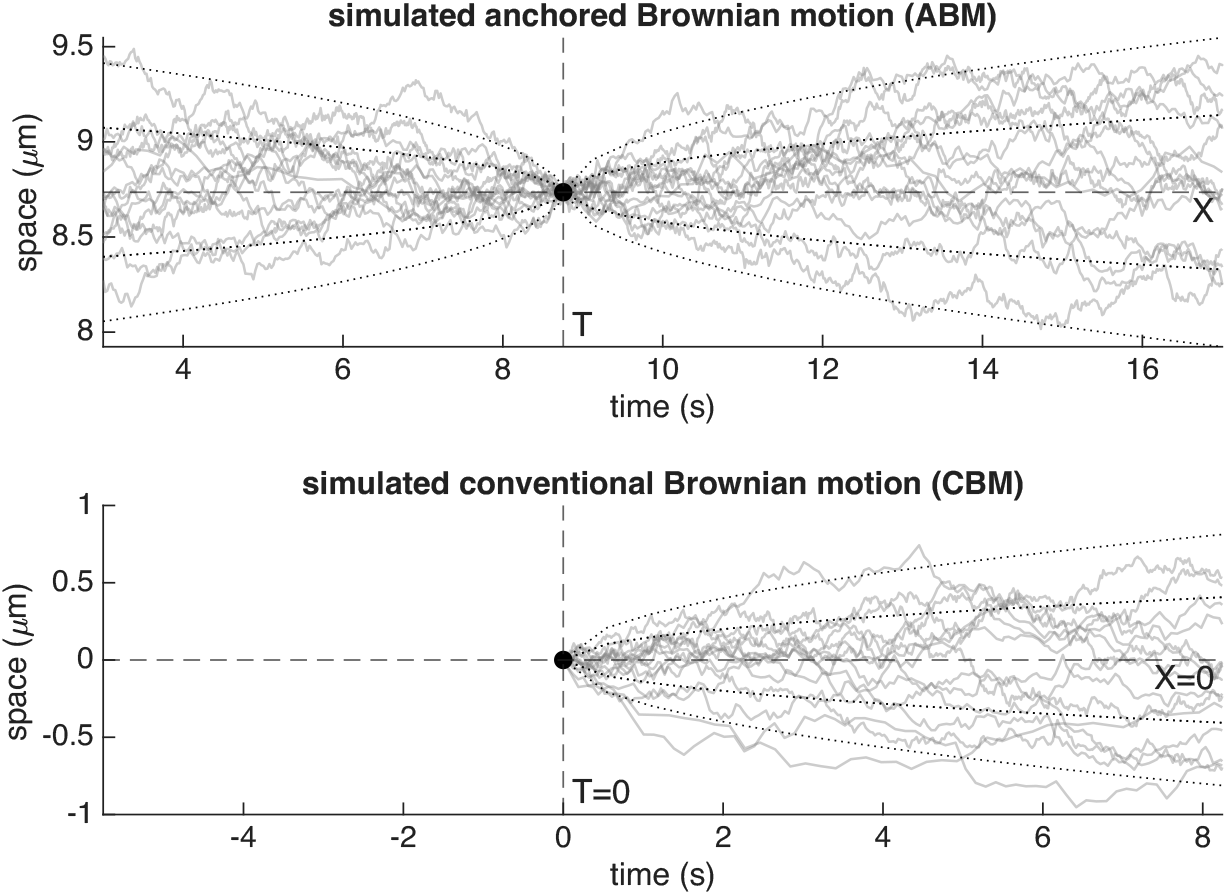
Comparison between ABM and CBM: The upper panel shows simulated ABM trajectories with given anchor and D = 0.01 µm^2^/s. The trajectories are generated with a fine time grid as described in section 2.2.1. The dashed lines indicate the positioning of the anchor and dotted lines indicate 1 and 2 standard deviations away from the position’s mean, ie 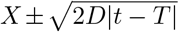, in accordance with eq. (2). The lower panel shows simulated CBM trajectories of the same D.

